# Repurposing Nintedanib for Pathological Cardiac Remodeling and Dysfunction

**DOI:** 10.1101/2020.12.21.423817

**Authors:** Prachi Umbarkar, Anand P. Singh, Sultan Tousif, Qinkun Zhang, Palaniappan Sethu, Hind Lal

## Abstract

**Background:** Heart Failure (HF) is the leading cause of death worldwide. Myocardial fibrosis, one of the clinical manifestations implicated in almost every form of heart disease, contributes significantly to HF development. However, there is no approved drug specifically designed to target cardiac fibrosis. Nintedanib (NTB) is an FDA approved tyrosine kinase inhibitor for idiopathic pulmonary fibrosis (IPF) and chronic fibrosing interstitial lung diseases (ILD). The favorable clinical outcome of NTB in IPF patients is well established. Furthermore, NTB is well tolerated in IPF patients irrespective of cardiovascular comorbidities. However, there is a lack of direct evidence to support the therapeutic efficacy and safety of NTB in cardiac diseases.

**Methods and Results:** We examined the effects of NTB treatment on cardiac fibrosis and dysfunction using a murine model of HF. Specifically, 10 weeks old C57BL/6J male mice were subjected to Transverse Aortic Constriction (TAC) surgery. NTB was administered once daily by oral gavage (50mg/kg) till 16 weeks post-TAC. Cardiac function was monitored by serial echocardiography. Histological analysis and morphometric studies were performed at 16 weeks post-TAC. In the control group, systolic dysfunction started developing from 4 weeks post-surgery and progressed till 16 weeks. However, NTB treatment prevented TAC-induced cardiac functional decline. In another experiment, NTB treatment was stopped at 8 weeks, and animals were followed till 16 weeks post-TAC. Surprisingly, NTB’s beneficial effect on cardiac function was maintained even after treatment interruption. NTB treatment remarkably reduced cardiac fibrosis as confirmed by Masson’s trichome staining and decreased expression of collagen genes (COL1A1, COL3A1). Compared to TAC group, NTB treated mice showed lower HW/TL ratio and cardiomyocyte cross-sectional area. Our *in vitro* studies demonstrated that NTB prevents myofibroblast transformation, TGFβ1-induced SMAD3 phosphorylation, and production of fibrogenic proteins (Fibronectin-1). However, NTB significantly altered vital signaling pathways in both, isolated fibroblast and cardiomyocytes, suggesting that its biological effect and underlying cardiac protection mechanisms are not limited to fibroblast and fibrosis alone.

**Conclusion:** Our findings provide a proof of concept for repurposing NTB to combat adverse myocardial fibrosis and encourage the need for further validation in large animal models and subsequent clinical development for HF patients.

## Introduction

Nintedanib (BIBF 1120) was developed by Boehringer Ingelheim (BI) Pharmaceuticals, Inc. In 2014, the FDA approved its use for the treatment of Idiopathic Pulmonary Fibrosis (IPF). BI is marketing NTB under the brand name Ofev. Recently (October, 2019) it obtained FDA’s “breakthrough therapy” designation and approval for treatment of chronic fibrosing interstitial lung diseases (ILD). Moreover, NTB is used as a second-line treatment along with other medications to treat non-small-cell lung cancer ^1^, and in several phases I to III clinical trials for different types of cancer (ClinicalTrials.gov Identifier: NCT02149108, NCT02902484, NCT03377023, NCT02399215). Currently, BI is recruiting patients in phase 3 clinical trial named NINTECOR (ClinicalTrials.gov Identifier: NCT04541680) to assess whether NTB slows the progression of lung fibrosis in COVID-19 survivors.

NTB is a potent tyrosine kinase inhibitor that primarily targets RTKs such as vascular endothelial growth factor receptor (VEGFR) 1, 2, and 3, platelet-derived growth factor receptor (PDGFR) α and β, and fibroblast growth factor receptor (FGFR) 1, 2, and 3. It also inhibits some non-RTKs including Lck, Lyn, and Src.^2^ Clinical trials revealed that NTB prevented the progression of fibrosis and improved lung function, leading to a better prognosis in IPF patients.^3^ Many pre-clinical studies have demonstrated that NTB has antifibrotic effects in disease models of different organs, including lung, liver, skin, and kidney. ^4-7^ Since the fundamental mechanisms of fibrosis remain common irrespective of etiology of disease, it is conceivable that NTB may also prevent adverse fibrotic remodeling implicated in cardiac diseases.

Fibrosis, characterized by excessive accumulation of ECM, is associated with multiple cardiac disorders. ECM deposition causes stiffening of the cardiac muscles, increases the risk of arrhythmia, disrupts electrical and mechanical coupling of cardiac cells, subsequently leading to functional decline and HF. Fibroblasts are the key players in ECM production and fibrosis progression.^8^ Several recent studies with FB-specific mouse models have demonstrated that activated fibroblasts and myocardial fibrosis contribute significantly to the HF. ^9-14^ However, at present, there is no therapy specifically designed to target myocardial fibrosis. In the current study, we determined the possibility of repurposing NTB to treat myocardial fibrosis and cardiac dysfunction. We employed a mouse model of pressure overload and examined the effect of NTB intervention on pressure overload-induced cardiac pathophysiology. We found that NTB, when administered as a preventive regimen, remarkably reduces pressure overload-induced cardiac fibrosis, hypertrophy, and dysfunction. Surprisingly these cardioprotective effects were observed despite interruption of the NTB treatment. Furthermore, we demonstrated that NTB alters fundamental signaling pathways in cardiac fibroblasts and cardiomyocytes, indicating multi-target mechanisms of action in the heart.

## Materials and Methods

### Mice and Nintedanib treatment

C57BL/6J mice were obtained from The Jackson Laboratories (Stock No: 000664), and the colony was maintained at the University of Alabama at Birmingham (UAB) animal care facility. All animal studies were approved by the Institutional Animal Care and Use Committee at UAB. At 10 weeks of age, mice were subjected to TAC surgery. NTB (Cat. No. N9055, LC Laboratories) was dissolved in water by heating to 50°C while intermittent stirring. NTB was administered daily by oral gavage (50mg/kg). NTB administration was started 1 day before surgery and continued as per experimental design.

### Transverse Aortic Constriction (TAC) surgery in mice

TAC was performed as previously described.^15^ Briefly, mice were sedated with isoflurane (induction, 3%; maintenance, 1.5%) and anesthetized to a surgical plane with intraperitoneal ketamine (50 mg/kg) and xylazine (2.5 mg/kg). Anesthetized mice were intubated, and a midline cervical incision was made to expose the trachea and carotid arteries. A blunt 20-gauge needle was inserted into the trachea and connected to a volume-cycled rodent ventilator on supplemental oxygen at a rate of 1 l/min, with a respiratory rate of 140 breaths/min. Aortic constriction was performed by tying a 7-0 nylon suture ligature against a 27-gauge needle. The needle was then promptly removed to yield a constriction of approximately 0.4 mm in diameter. To confirm the efficiency of consistent TAC surgery, the pressure gradient across the aortic constriction was measured by Doppler echocardiography.

### Echocardiography

Echocardiography was performed as described previously.^16^ In brief, transthoracic M-mode echocardiography was performed with a 12-MHz probe (VisualSonics) on mice anesthetized by inhalation of isoflurane (1-1.5%). LV end-systolic interior dimension (LVID; s), LV end-diastolic interior dimension (LVID;d), LV end-diastolic posterior wall thickness (LVPW, d), LV end-systolic posterior wall thickness (LVPW, s), ejection fraction (EF), and fractional shortening (FS) values were obtained by analyzing data using the Vevo 3100 program.

### Histology

Heart tissues were fixed in 10% neutral buffered formalin, embedded in paraffin, and sectioned at 5 μm thickness. Sections were stained with Masson Trichrome (HT15, Sigma-Aldrich) as per the manufacturer’s instruction. The images of the LV region were captured using a Nikon eclipse E200 microscope with NIS element software version 5.20.02. The quantification of LV fibrosis and cross-sectional area of cardiomyocytes (CSA) were determined with ImageJ version 1.52a software (NIH).

### Western blot

LV tissues were dissected from mouse heart and homogenized with cell lysis buffer (Cell signaling Technology) having 50 mM Tris-HCl (pH7.4), 150 mM NaCl, 1mM EDTA, 0.25% sodium deoxycholate, 1% NP-40, and freshly supplemented Protease Inhibitor Cocktail (Sigma-Aldrich) and Phosphatase Inhibitor Cocktail (Sigma-Aldrich). Protein concentration was determined with the Bradford assay (Bio-Rad protein assay kit). An equal amount of proteins was denatured in SDS–PAGE sample buffer, resolved by SDS–PAGE and transferred to Immobilon-P PVDF membrane (EMD Millipore). The membranes were blocked in Odyssey blocking buffer (LI-COR Biosciences) for 1h at RT. Primary antibody incubations were performed at different dilutions as described in Supplemental Table 1. All incubations for primary antibodies were done overnight at 4°C and followed by secondary antibody (IRDye 680RD or IRDye 800CW from LI-COR Biosciences) incubation at 1:3000 dilutions for 1h at RT. Proteins were visualized with the Odyssey Infrared Imaging System (LI-COR Biosciences). Band intensity was quantified by Image Studio version 5.2 software.

### RNA extraction and quantitative PCR analysis

Total RNA was extracted from heart tissue using the RNeasy Mini Kit (74104, Qiagen) according to the manufacturer’s protocol. cDNA was synthesized using the iScript cDNA synthesis kit (170-8891, Bio-Rad) following the manufacturer’s instructions. Gene expression was analyzed by quantitative PCR (qPCR) using the TaqMan Gene Expression Master Mix (4369016, Applied Biosystems) and TaqMan gene expression assays (Applied Biosystems) on a Quant studio 3 (Applied Biosystems) Real-Time PCR Detection machine. Details of TaqMan gene expression assays used in this study are provided as Supplemental Table 2. Relative gene expression was determined by using the comparative CT method (2^-ΔΔCT^) and was represented as fold change. Briefly, the first ΔCT is the difference in threshold cycle between the target and reference genes: ΔCT= CT (a target gene X) − CT (18S rRNA) while ΔΔCT is the difference in ΔCT as described in the above formula between the CTL and KO group, which is = ΔCT (KO target gene X) − ΔCT (CTL target gene X). Fold change is calculated using the 2-ΔΔCT equation.^17^

### Isolation and culture of neonatal rat cardiomyocytes and fibroblasts

Primary cultures of neonatal rat ventricular cardiomyocytes (NRVM) and fibroblasts (NRVF) were prepared from 1-2-day-old pups of Sprague-Dawley rats as described previously.^18^

### Cell migration assay

NRVFs were seeded in culture dishes and allowed to grow till confluency. The wound was created by scratching the monolayer of the cells. Cells were pre-treated with NTB (0.5µM) for 1h followed by agonists treatments for a total of 24h. The percentage of wound closure was monitored and quantified over the indicated time points.

### Statistical analysis

Analyses were performed using GraphPad Prism (version 9.0.0). Differences between more than 2 data groups were evaluated for significance by ANOVA followed by Tukey’s multiple comparison test. All data are expressed as mean ± SEM. For all tests, a p-value of < 0.05 was considered statistically significant.

## Results

### NTB administration prevented pressure overload-induced cardiac dysfunction and adverse cardiac remodeling

We evaluated the therapeutic efficacy of NTB in a pre-clinical mouse model of heart failure. We subjected 10 weeks old C57BL/6J mice to TAC surgery. To study the effect of nintedanib on pressure overload-induced cardiac pathology, the drug was administered throughout the experimental period (preventive regimen) **(Figure 1A)**. We examined cardiac function by serial M-mode echocardiography. In the TAC group, LVEF and LVFS started to decline from 4 weeks post-surgery, indicating systolic dysfunction **(Figure 1B-1C)**. These changes were associated with the development of structural remodeling as reflected by a significant increase in LVIDs, LVPWDs, and LV Mass in the TAC group. All these parameters were remarkably normalized due to NTB administration, indicating NTB treatment improved cardiac function and prevented adverse cardiac remodeling in TAC mice **(Figure 1D-1H)**.

**Figure 1:**
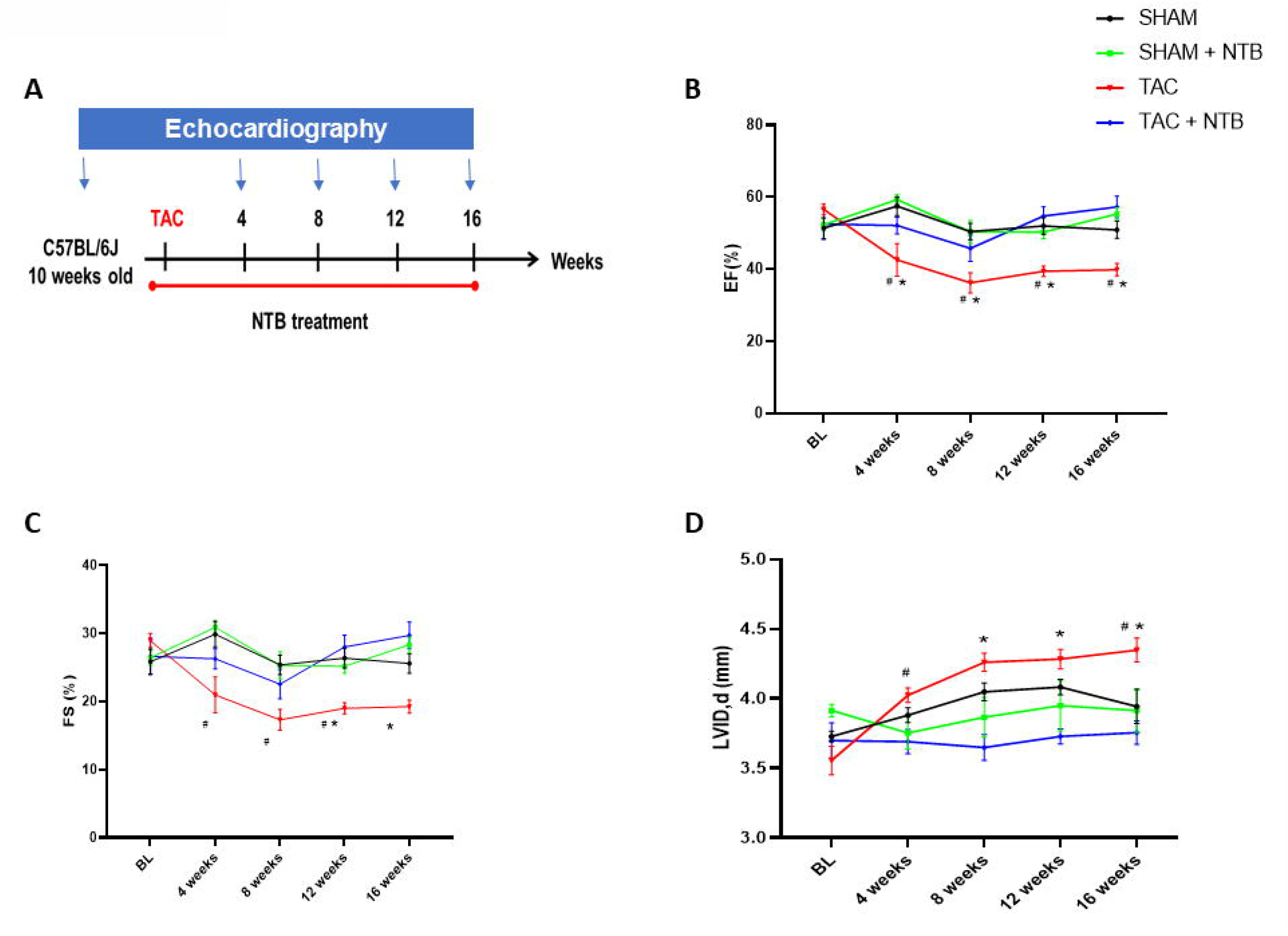
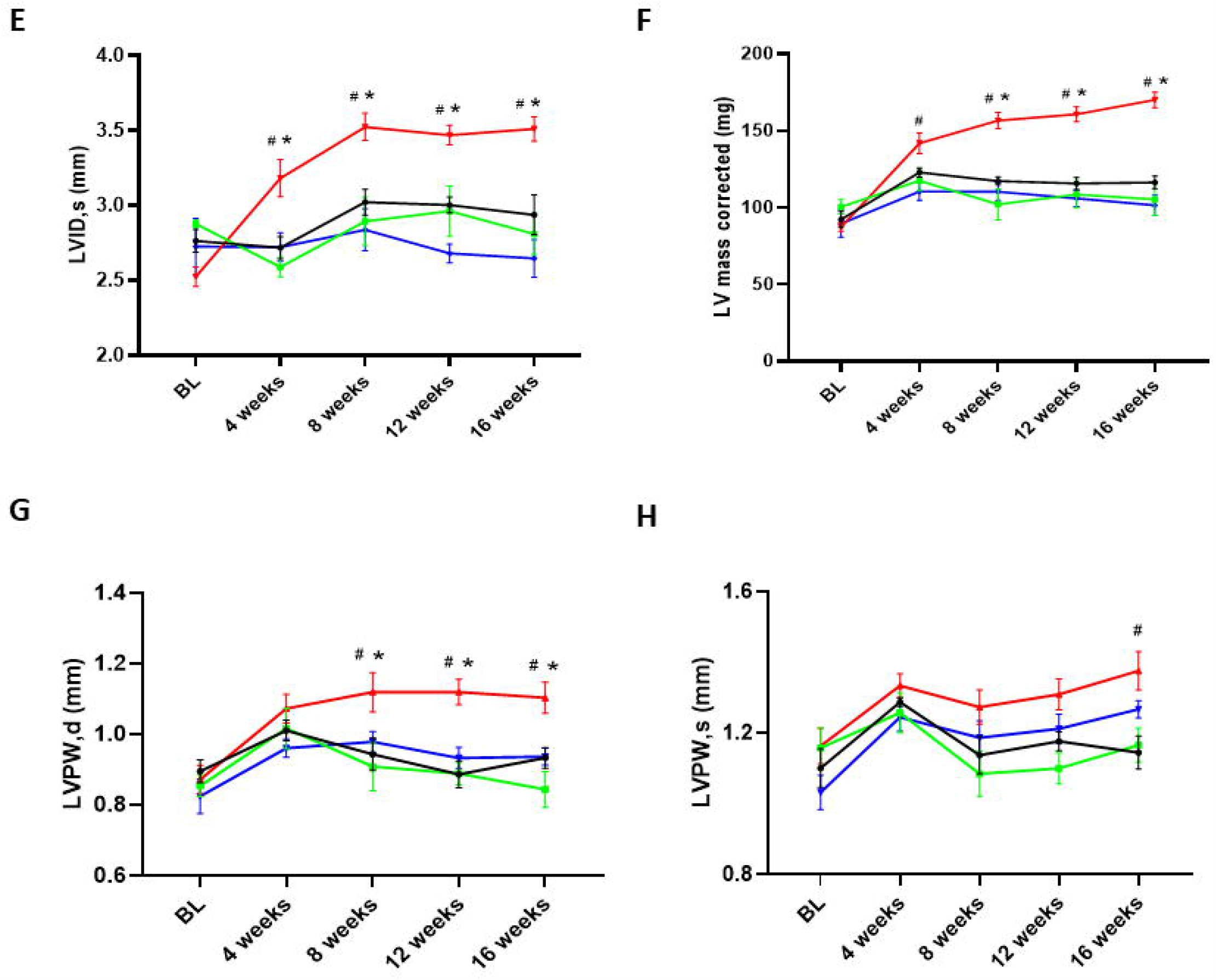
NTB treatment improves cardiac function in TAC mice. (A) At 10 weeks of age, C57BL/6J mice were subjected to TAC surgery and treated with NTB (50mg/kg/day) for the indicated time. Cardiac function was assessed by serial echocardiography (B) EF: Ejection fraction, (C) FS: Fractional shortening, (D) LVID, d: LV end-diastolic dimension (E) LVID, s: LV end-systolic dimension, (F) LV Mass corrected (G) LVPW, d: LV end-diastolic posterior wall thickness (H) LVPW, s: LV end-systolic posterior wall thickness. (n>5 per group) Data are mean ± SEM. ^#^*P*<0.05 for SHAM vs TAC, \**P*<0.05 for TAC vs.TAC + NTB.

To verify the effects of NTB treatment on adverse cardiac remodeling, we examined cardiac hypertrophy and fibrosis at 16 weeks post-TAC. We measured HW/TL ratio and CSA. Both these parameters were elevated in the TAC group revealing hypertrophic remodeling **(Figure 2A & 2B)**. For assessment of cardiac fibrosis, we did Masson’s Trichome staining and found excessive collagen deposition in TAC hearts **(Figure 2C & 2D)**. All these characteristics of pathological cardiac remodeling were alleviated in NTB treated TAC group. Additionally, we compared expression levels of key genes related to cardiac fibrosis (COL1A1 and COL3A1). These molecular markers were significantly augmented in the TAC group and are normalized in NTB treated TAC mice **(Figure 2E & 2F)**.

**Figure 2:**
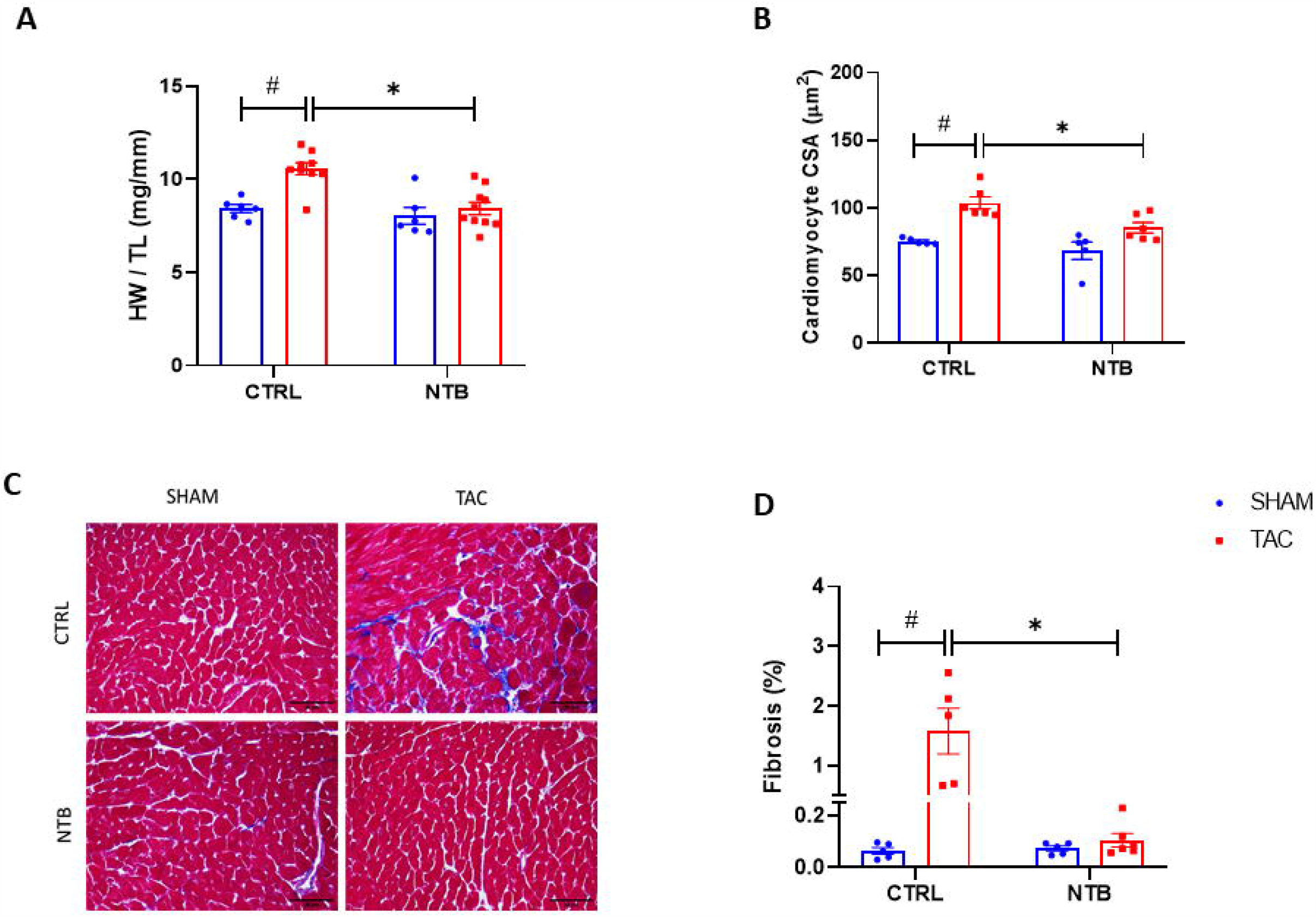
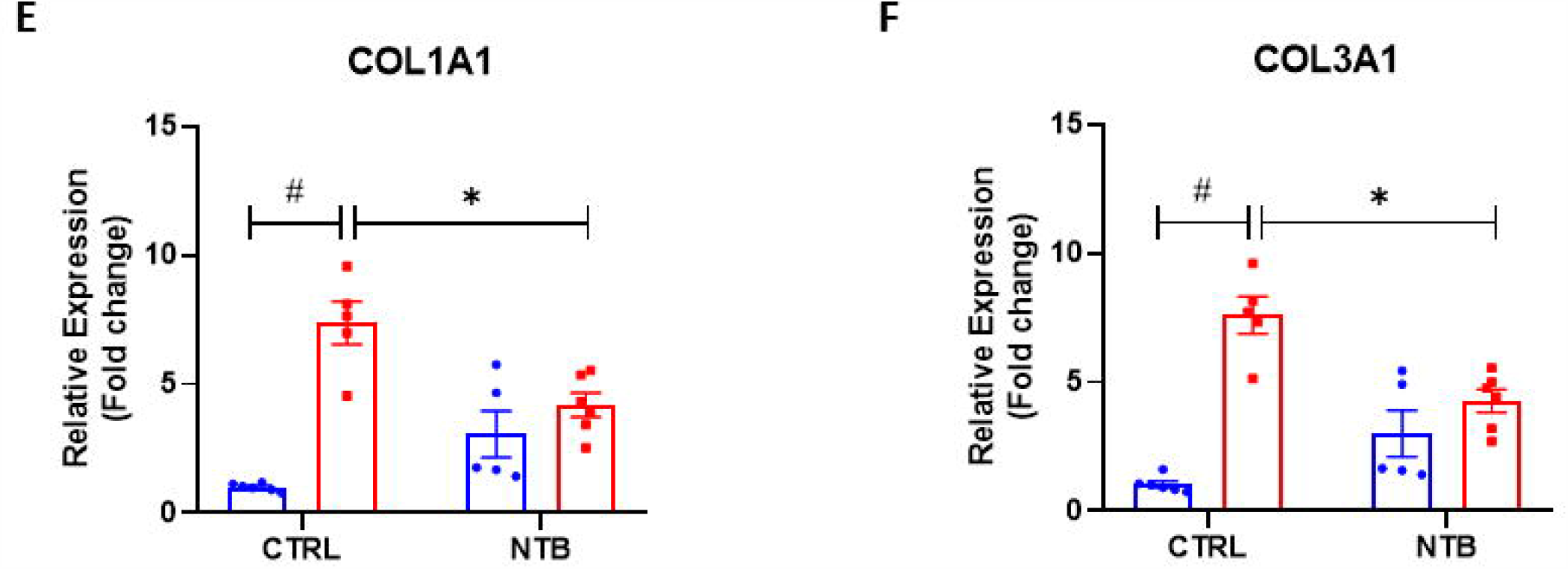
NTB treatment prevents pressure overload induced cardiac fibrosis and hypertrophy. Morphometric studies were performed at 16 weeks after TAC surgery; (A) Heart weight (HW) to tibia length (TL) ratio, (B) Quantification of cardiomyocyte cross-sectional area (CSA), (C) Masson’s trichome staining. Representative trichrome-stained LV regions and (D) Quantification of LV fibrosis. Scale bar = 30 µm. RNA was extracted from left ventricle of experimental animals and gene expression analysis was carried out by qPCR (E) COL1A1 (F) COL3A1. (n>5 per group) Data are mean ± SEM. ^#^*P*<0.05 for SHAM vs. TAC, \**P*<0.05 for TAC vs. TAC + NTB.

### Cardioprotective effects of NTB persisted despite interruption of the treatment

To test whether the efficacy of NTB is maintained even after cessation of the treatment, we treated mice with NTB for 8 weeks post-TAC, thereafter drug administration was discontinued, and animals were followed till 16 weeks post-TAC **(Figure 3A)**. We assessed cardiac function by serial echocardiography. We found that cardiac function started to deteriorate from 4 weeks post-surgery in TAC groups. When compared to SHAM, LVIDs, LVPWDs, and LV mass were significantly increased in TAC groups indicating the development of adverse cardiac remodeling. As observed in the preventive treatment regimen, continuous NTB treatment till 8 weeks prevented TAC induced cardiac dysfunction and remodeling. Surprisingly, these cardioprotective effects of NTB administration were sustained up even after 8 weeks of washout period **(Figure 3B-3H)**.

**Figure 3:**
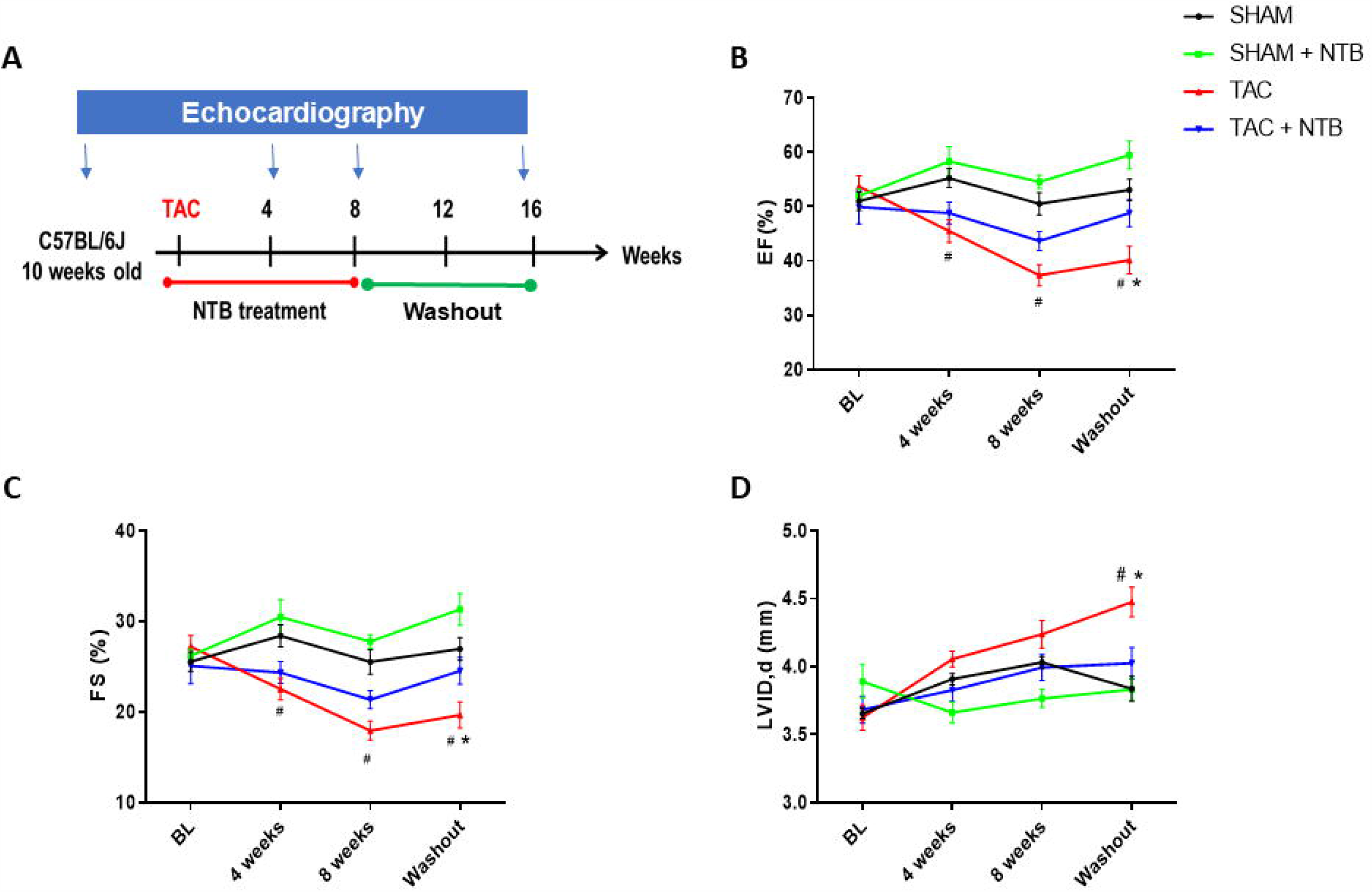
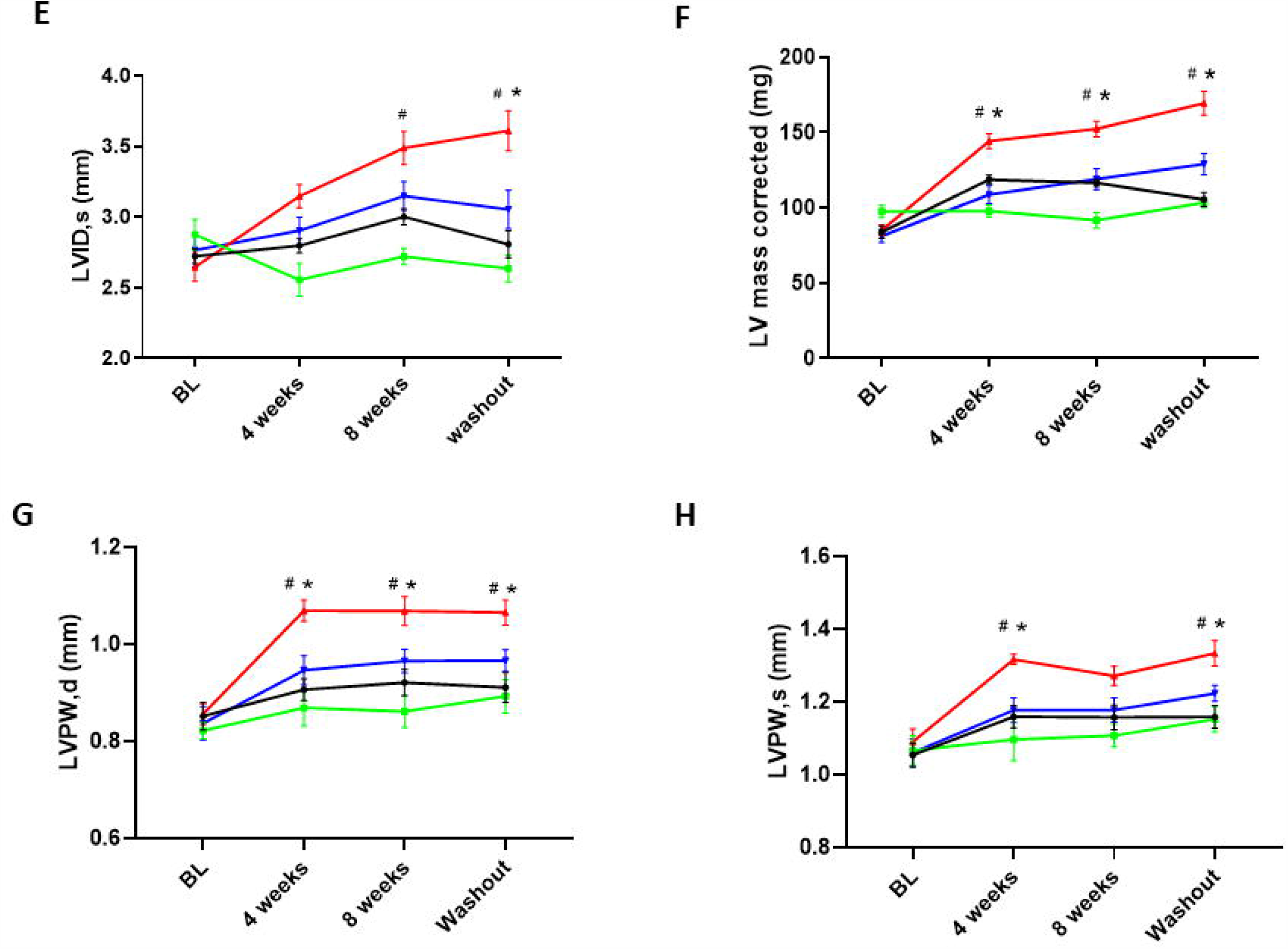
Interruption of the treatment does not affect cardiac function improvement in NTB treated TAC mice. (A) At 10 weeks of age, C57BL/6J mice were subjected to TAC surgery and treated with NTB (50mg/kg/day) for 8 weeks. After 8 weeks, NTB treatment was discontinued, and animals were followed for additional 8 weeks. Cardiac function was assessed by serial echocardiography (B) EF: Ejection fraction, (C) FS: Fractional shortening, (D) LVID, d: LV end-diastolic dimension (E) LVID, s: LV end-systolic dimension, (F) LV Mass corrected, (G) LVPW, d: LV end-diastolic posterior wall thickness, and (H) LVPW, s: LV end-systolic posterior wall thickness (n>5 per group) Data are mean ± SEM. ^#^*P*<0.05 for SHAM vs. TAC, \**P*<0.05 for TAC vs. TAC + NTB.

To further verify the cardioprotective effects of NTB treatment on cardiac remodeling, we harvested the heart of experimental animals at 8 weeks washout period for morphometrics and histological studies. TAC group showed cardiac hypertrophy as reflected by a significant increase in HW/TL and CSA compared with the SHAM group **(Figure 4A & 4B)**. Masson’s Trichome staining revealed excessive collagen deposition in TAC groups **(Figure 4C & 4D)**. Importantly, cardiac hypertrophy and fibrosis were reduced in mice treated with NTB. Next, we compared the expression of genes related to cardiac fibrosis (COL1A1, COL3A1) using the qPCR method. In the TAC group, there was a significant increase in levels of these molecular markers and NTB treatment could normalize their levels **(Figure 4E & 4F)**. All these results indicate the maintained efficacy of NTB despite the interruption of the treatment.

**Figure 4:**
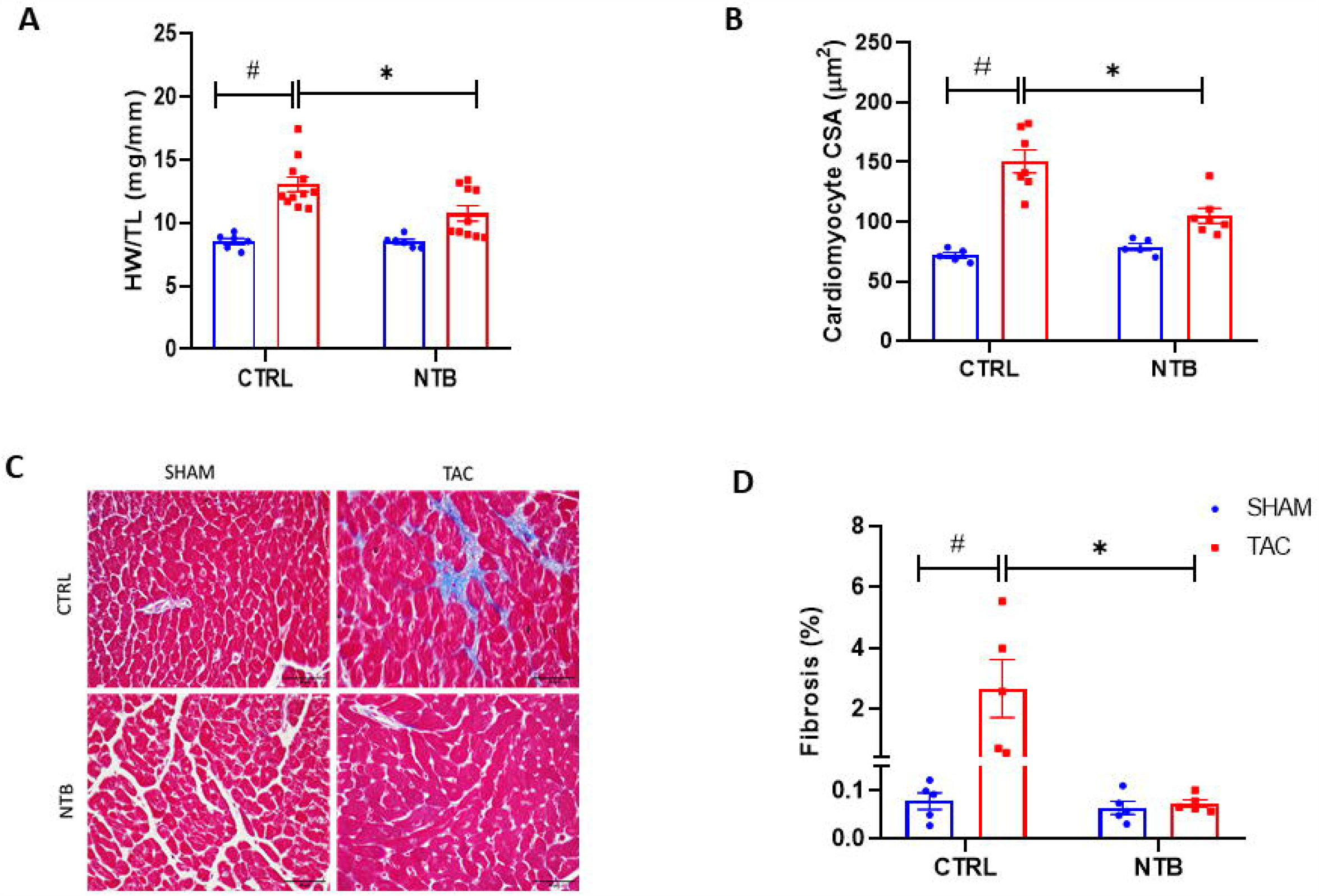
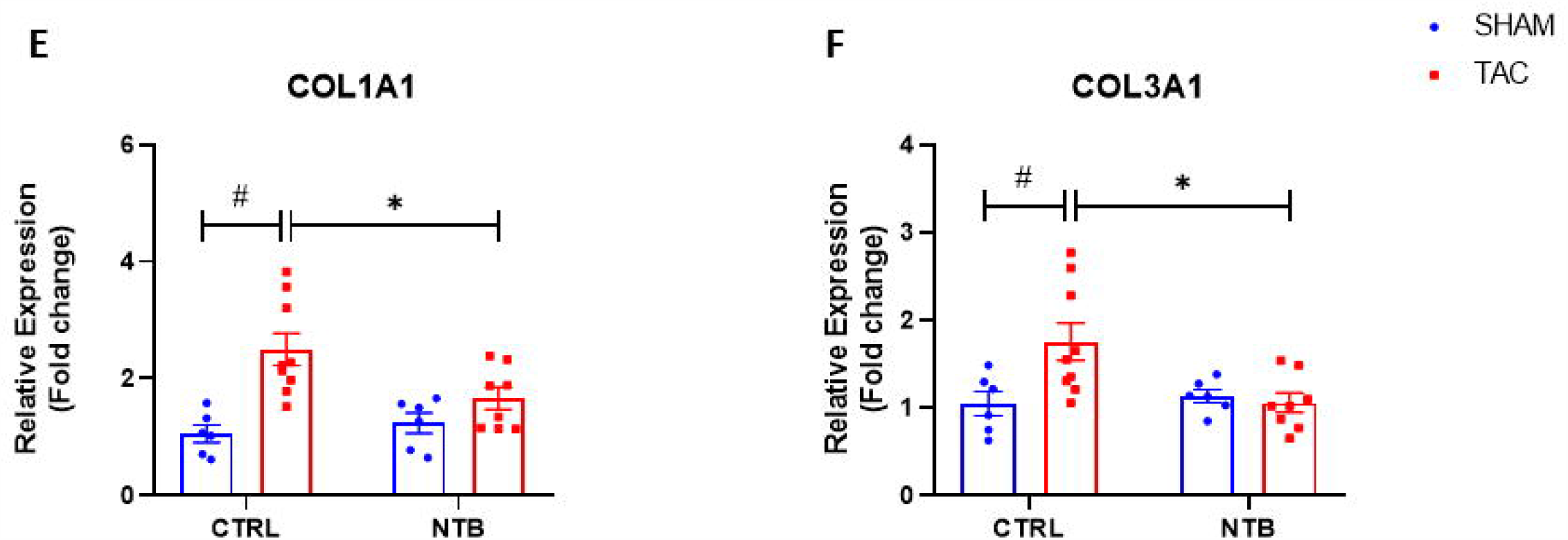
Effect of NTB treatment interruption on pressure overload induced adverse cardiac remodeling. At 10 weeks of age, C57BL/6J mice were subject to TAC surgery and treated with NTB (50mg/kg/day) for 8 weeks. After 8 weeks, NTB treatment was discontinued for further 8 weeks. Morphometric studies were performed at the end of washout period; (A) Heart weight (HW) to tibia length (TL) ratio, (B) Quantification of cardiomyocyte cross-sectional area (CSA), (C) Masson’s trichome staining. Representative trichrome-stained LV regions and (D) Quantification of LV fibrosis. Scale bar = 30 µm. RNA was extracted from left ventricle of experimental animals and gene expression analysis was carried out by qPCR (E) COL1A1 (F) COL3A1. (n>5 per group) Data are mean ± SEM. ^#^*P*<0.05 for SHAM vs. TAC, \**P*<0.05 for TAC vs. TAC + NTB.

### NTB treatment prevented profibrotic responses in cardiac fibroblasts

NTB has proven anti-fibrotic potential in disorders such as IPF. Since we observed a reduction in cardiac fibrosis in NTB treated TAC mice, we examined cardiac fibroblasts specific effects of NTB. Cardiac fibroblasts were pre-treated with NTB (0.5µM) for 1h followed by TGFβ1 treatment (10ng/ml for 1h). Our western blot results showed that NTB treatment prevented TGF-β1 mediated SMAD3 phosphorylation, a key pathway involved in myofibroblast transformation and fibrosis **(Figure 5A)**. NTB targets FGFR and inhibits RTKs associated with it. Since FGF/FGFR signaling is fundamental in fibroblast growth and survival, we studied the effects of NTB on basic fibroblast growth factor (bFGF) induced MAPK and AKT activation in cardiac fibroblasts. Cardiac fibroblasts were pre-treated with NTB (0.5µM) for 1h followed by bFGF treatment (25ng/ml bFGF for 10 min). As anticipated, NTB treatment prevented bFGF induced ERK and AKT activation in cardiac fibroblasts **(Figure 5B & 5C)**. However, NTB did not affect bFGF mediated P38 signaling **(Figure 5D)**. Additionally, NTB treatment inhibited agonists-induced cell migration and production of extracellular matrix protein such as fibronectin **(Figure 5E & 5F)**. These results indicate that NTB prevents myofibroblast transformation and fibrotic responses in cardiac fibroblasts.

**Figure 5:**
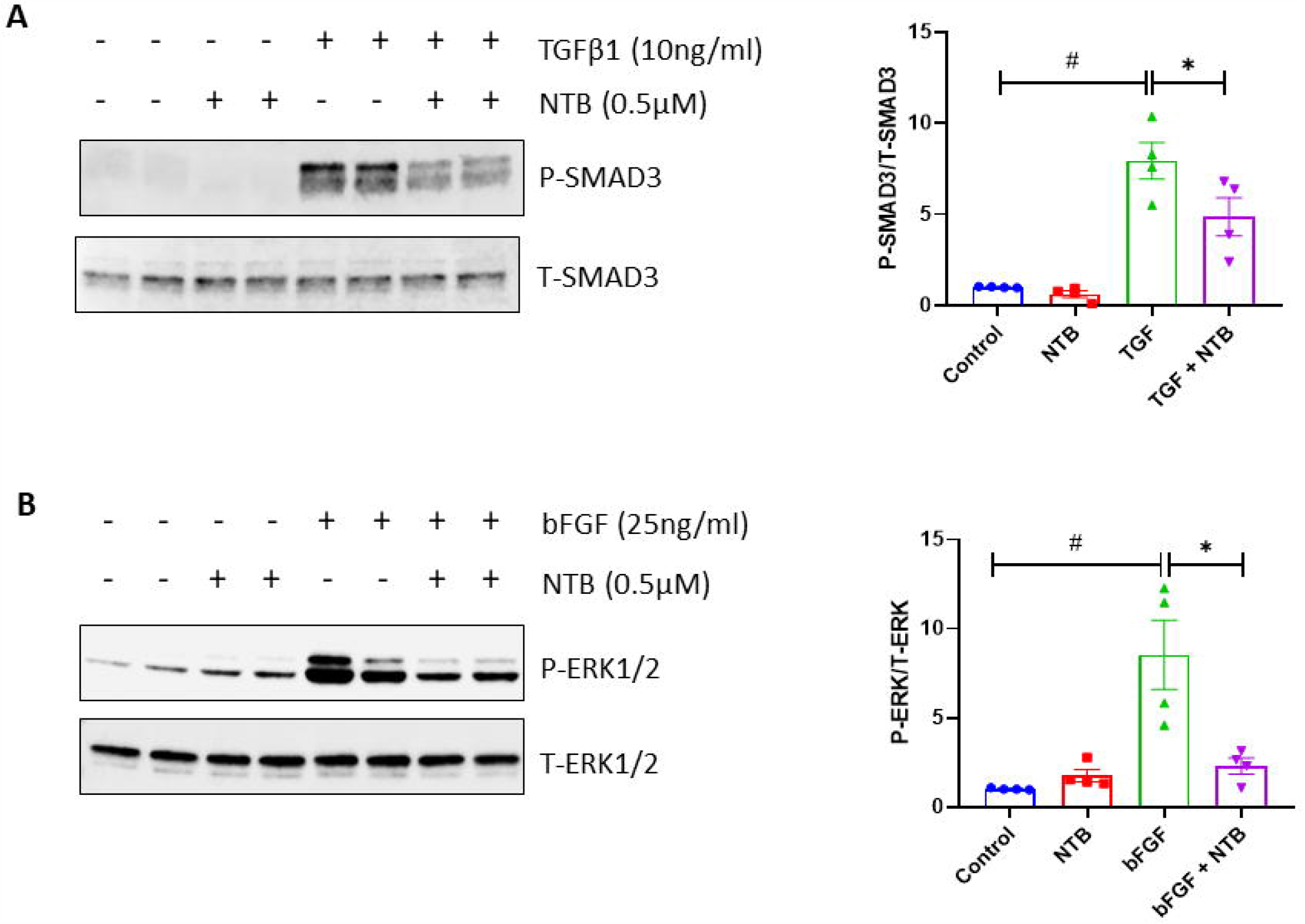
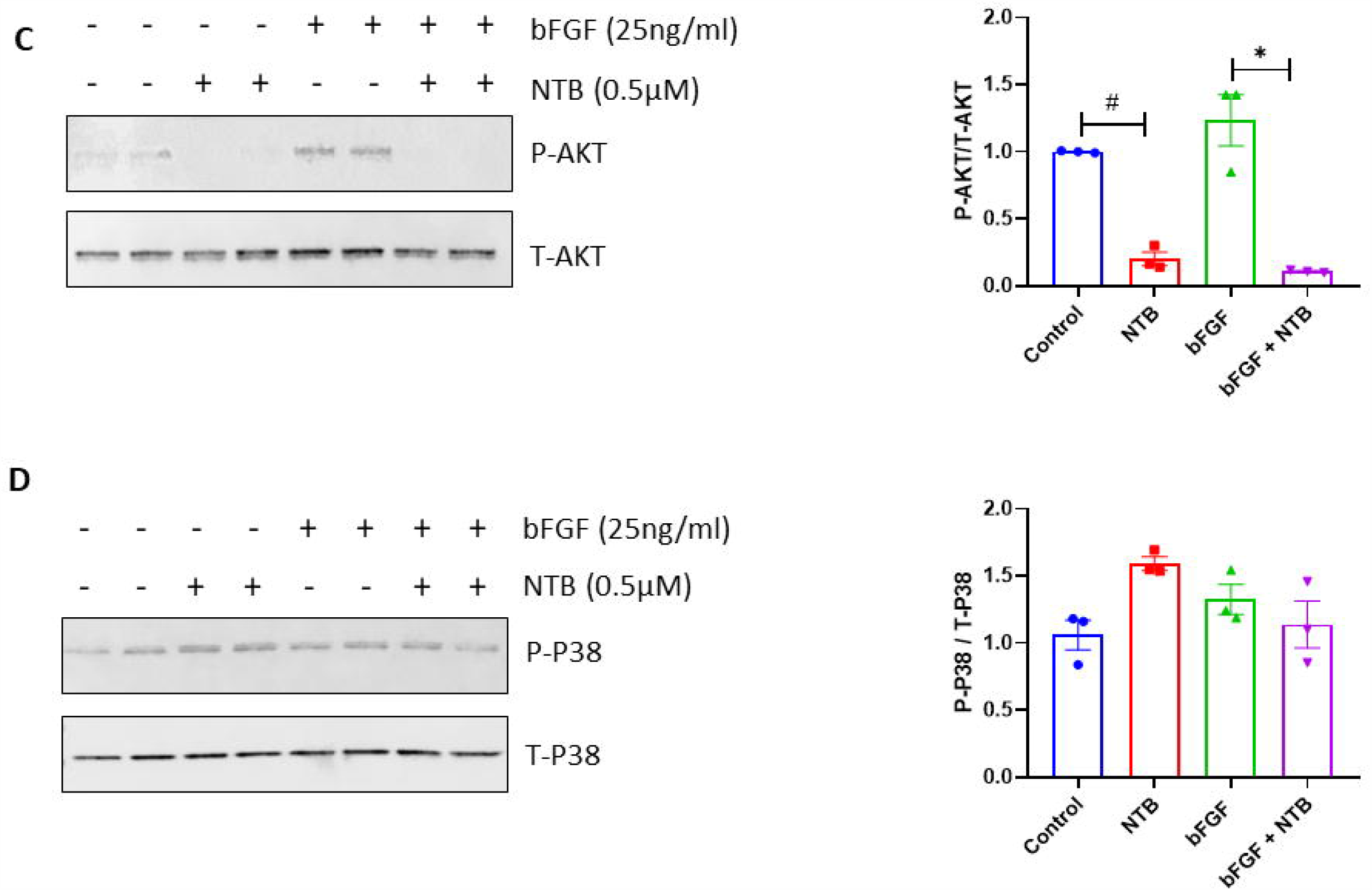
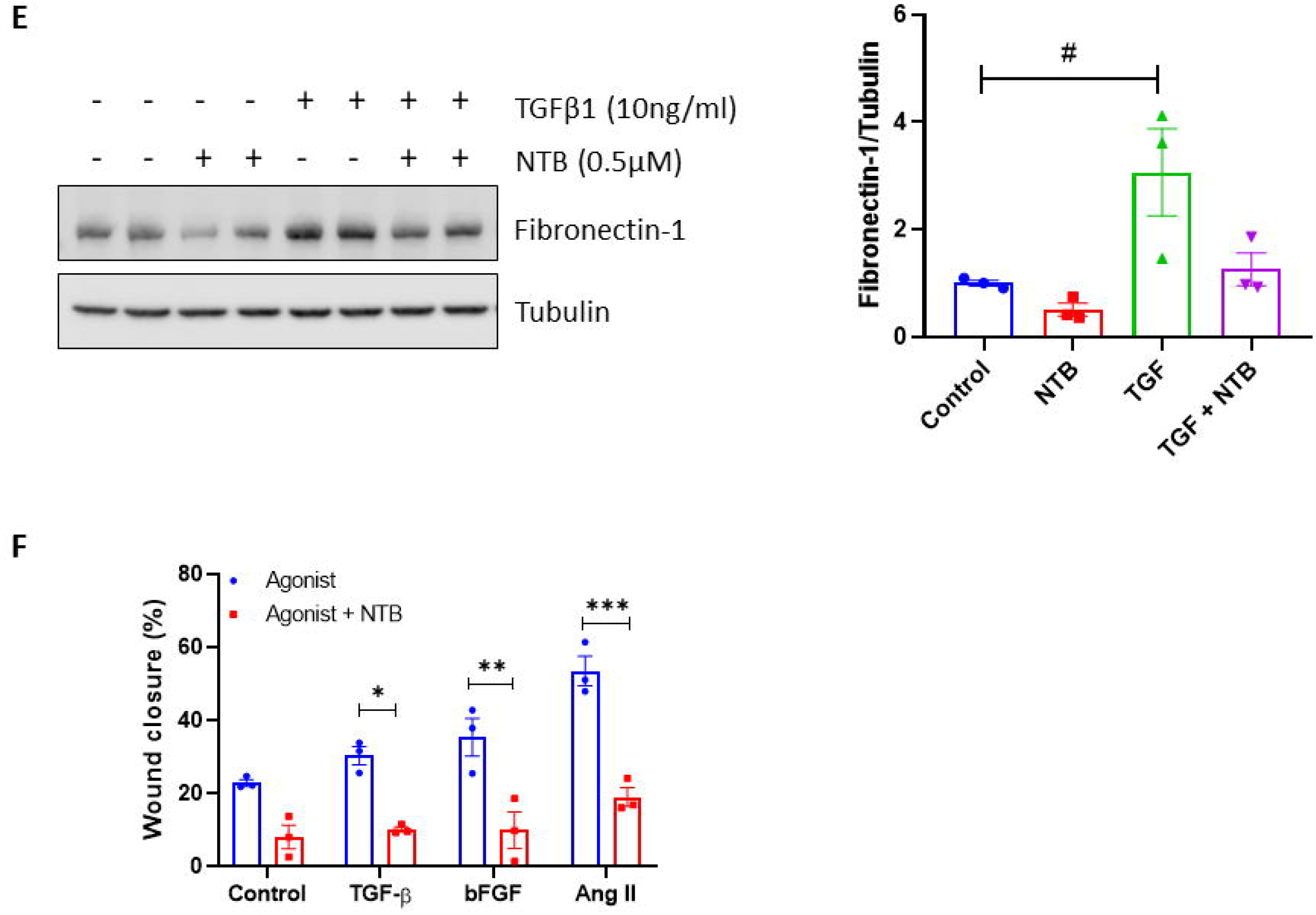
NTB treatment prevents profibrotic responses in cardiac fibroblasts. Neonatal rat cardiac fibroblasts (CFs) were pretreated with NTB (0.5µM) for 1h followed by agonists treatment (10ng/ml TGFβ1 for 1h or 25ng/ml bFGF for 10 min). Proteins were extracted, and Western blot analysis was carried out. Representative blots and Quantification - (A) SMAD3 (B) ERK (C) AKT (D) P38, and (E) Fibronectin-1. For cell migration assay, CFs were treated with agonists (10ng/ml TGFβ1, 25ng/ml bFGF, or 10µM AngII) in presence or absence of NTB (0.5µM) for 24h. Wound closure was monitored from 0h-24h and % wound closure was calculated at 24h using imageJ software (F). (n=3-4) Data are mean ± SEM. ^#^*P*<0.05 vs. Control, \**P*<0.05 vs. Agonist.

### Effect of NTB treatment on cardiomyocytes

Our *in vivo* studies showed that NTB treatment prevented pressure overload-induced cardiac hypertrophy. To evaluate whether the reduction in cardiac hypertrophy is due to the direct effect of NTB on cardiomyocytes (CM), we treated CMs with hypertrophy inducing agents [100µM phenylephrine (PE), 25ng/ml bFGF, or 10µM Angiotensin II (AngII)] in presence or absence of NTB (1µM) for 48h. Induction of hypertrophic response was confirmed by analyzing gene expression levels of ANP and BNP. We observed that NTB treatment had no significant effect on agonists induced hypertrophic response **(Figure 6A & 6B)**. Next, we verified the effect of NTB treatment on key signaling pathways in CMs. CMs were pre-treated with NTB (1µM) for 1h followed by phenylephrine (100µM) treatment for 15min. Western blot analysis showed that phenylephrinE-induced MAPK and AKT phosphorylation was inhibited by NTB (**Figure 6C-6F)**. Taken together, these results indicate that NTB can directly modulate CM signaling underlying hypertrophic stimuli. However, this much interference is not sufficient to prevent CM hypertrophic growth.

**Figure 6:**
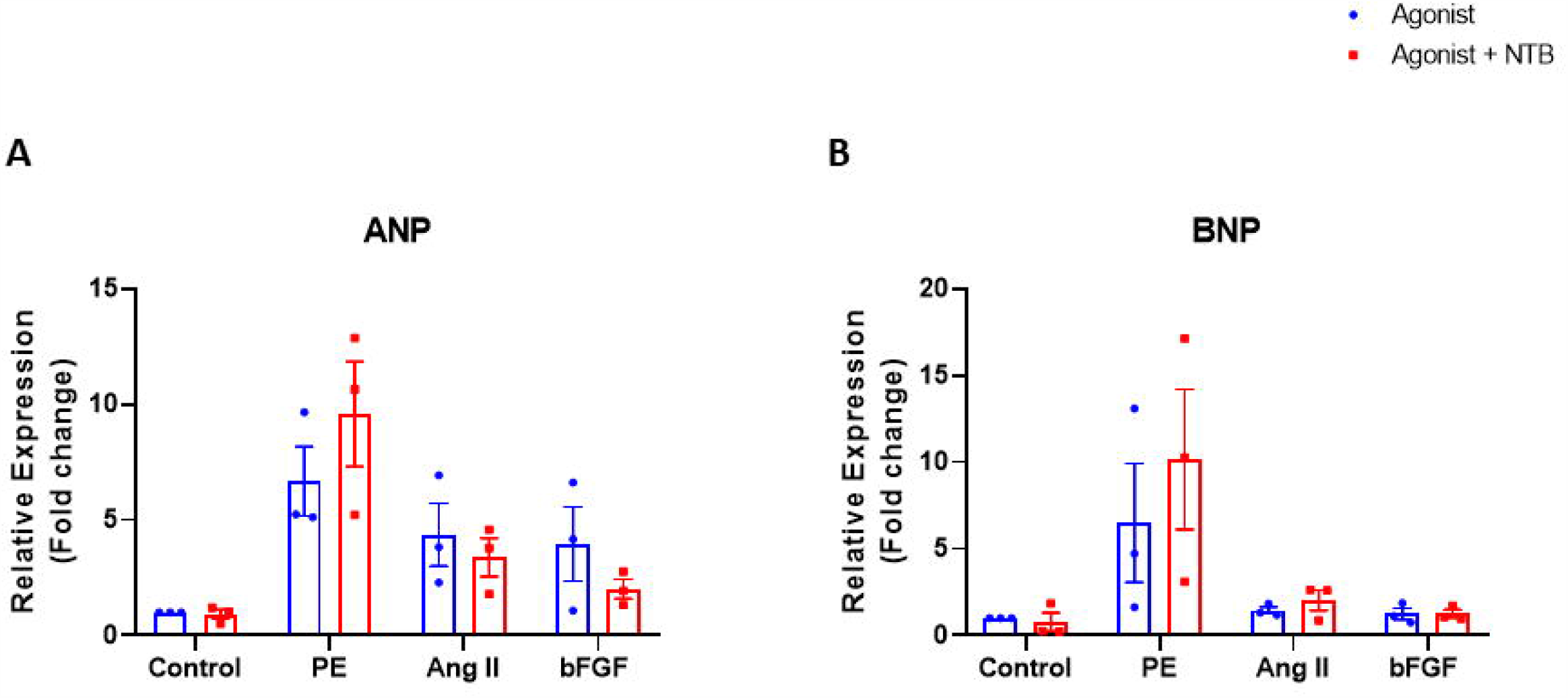
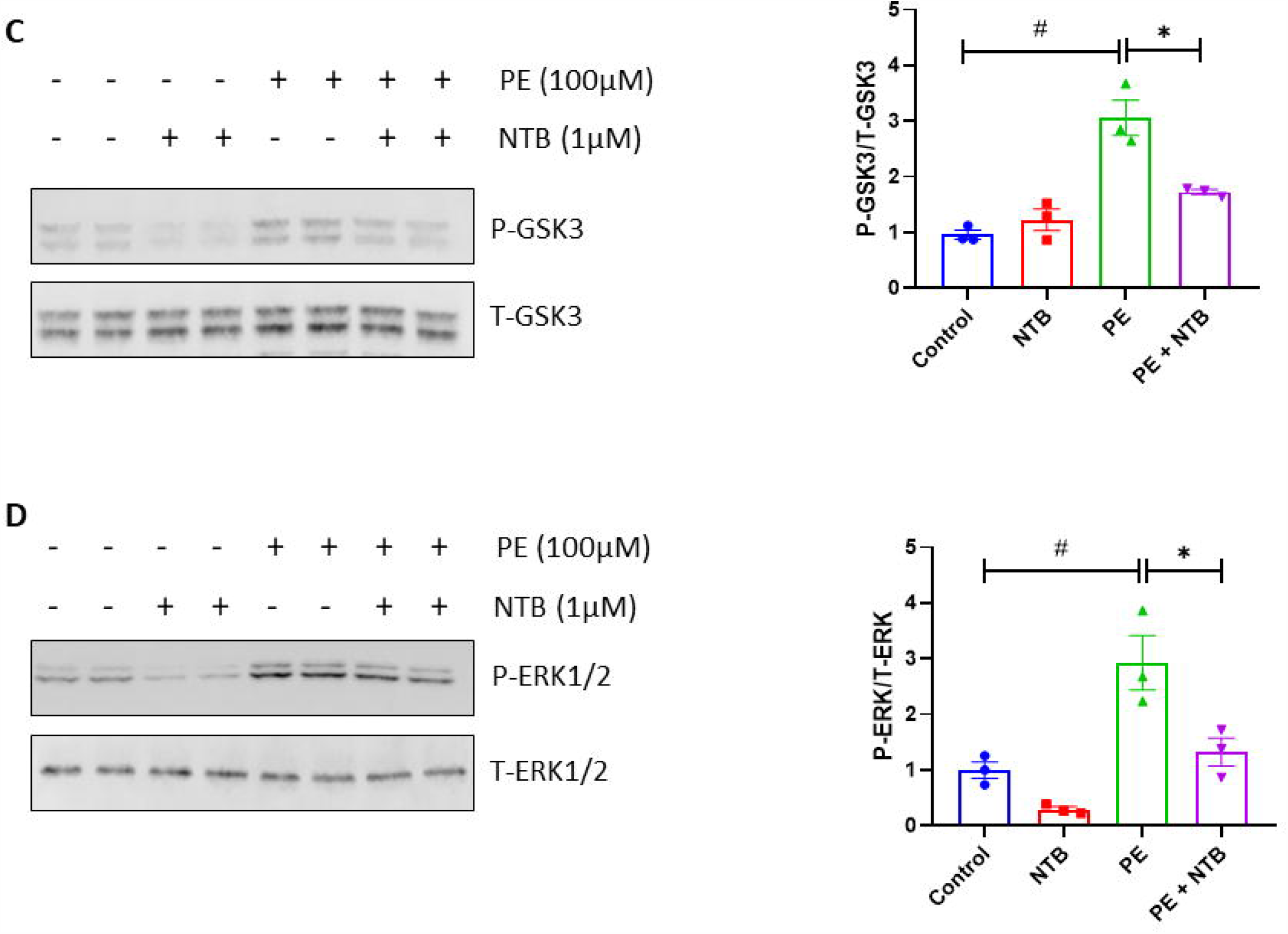
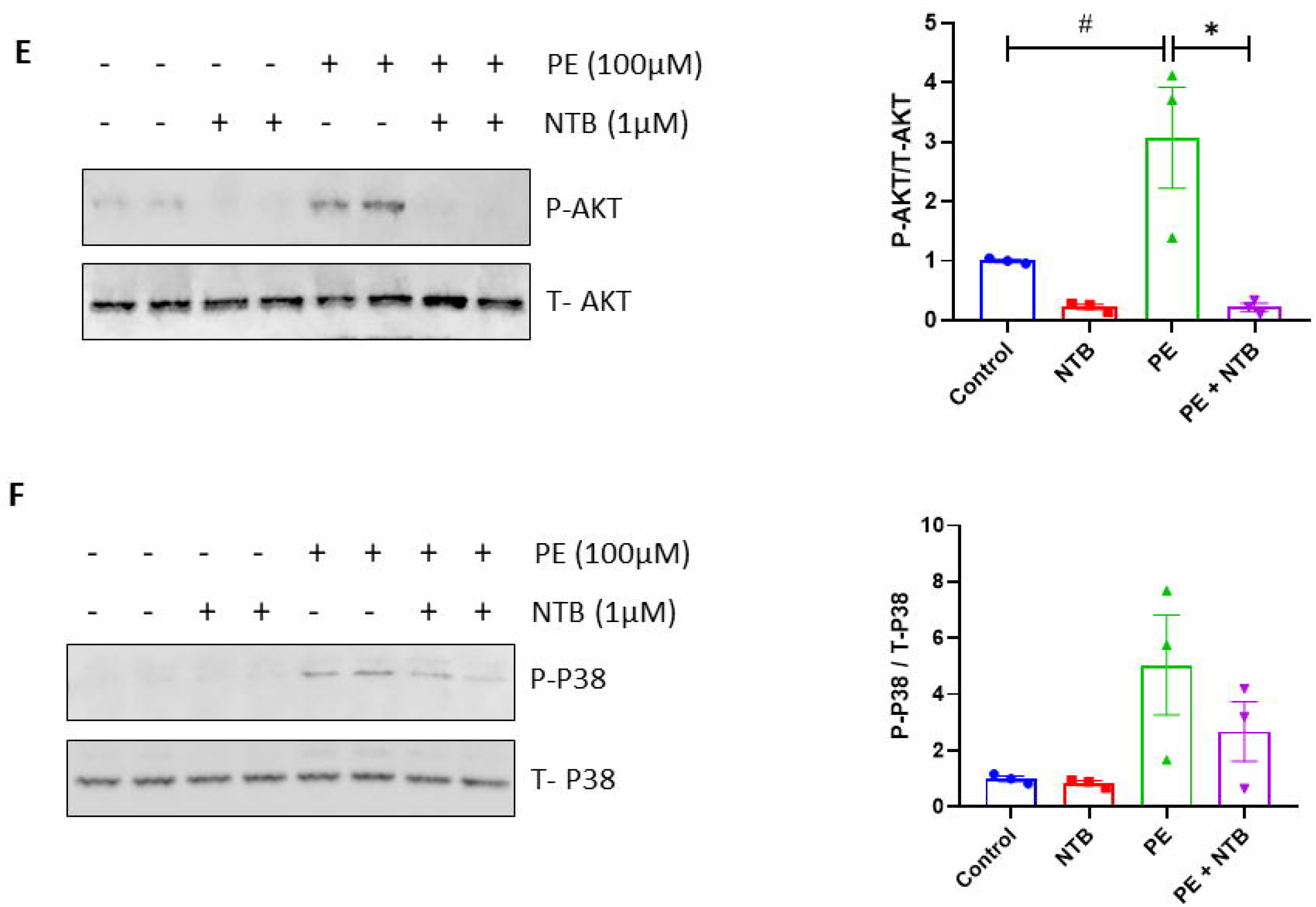
NTB treatment inhibits signaling pathways underlying hypertrophic stimuli in cardiomyocytes. To study whether NTB could prevent cardiomyocyte hypertrophy, neonatal rat cardiomyocytes (CMs) were treated with agonists (100µM PE, 25ng/ml bFGF, or 10µM AngII) in presence or absence of NTB (1µM) for 48h, RNA was extracted and gene expression levels were compared using qPCR method (A) ANP and (B) BNP. (n=3) Data are mean ± SEM. ^#^*P*<0.05 vs. Control, \**P*<0.05 vs. Agonist. To examine the effect of NTB on CM’s signaling, CMs were pretreated with NTB (1µM) for 1h followed by PE (100µM) treatment for 15min. Proteins were extracted, and Western blot analysis was carried out. Representative blots and Quantification - (C) GSK3α/β (D) ERK (E) AKT, and (F) P38. (n=3) Data are mean ± SEM. ^#^*P*<0.05 vs. Control, \**P*<0.05 vs. PE.

## Discussion

In the present study, we examined the effects of NTB treatment on pathological remodeling and cardiac dysfunction in a murine model of pressure overload. We found that when administered as a preventive regimen, NTB prevents pressure-overload induced myocardial dysfunction, adverse fibrotic remodeling, and pathological hypertrophy in mice. Importantly, the cardioprotective effects of NTB were sustained despite the cessation of the treatment.

NTB is approved for treatment in IPF patients, where its beneficial effects are attributed to its anti-fibrotic action and improvement of lung function.^3^ In the *mdx* mouse model of Duchenne Muscle Dystrophy (DMD), NTB administration reduced fibrosis and improved muscle performance.^19^ The efficacy of NTB has been tested in the pre-clinical model of Pulmonary arterial hypertension (PAH) in which authors demonstrated that NTB treatment reduced fibrosis and prevented RV hypertrophy.^20^ All these studies revealed that halting the progression of adverse fibrotic remodeling facilitates the functional improvement of the organ. In line with these reports, we observed that after NTB intervention, cardiac function is improved in the murine model of heart failure. This functional gain was associated with a significant decrease in cardiac fibrosis.

Nintedanib is a TKI that primarily targets PDGFR, VEGFR, and FGFR. There are multiple FDA approved TKIs with similar inhibition profiles. Most of these targets are associated with adverse cardiovascular events such as arrhythmia, PAH, and atherosclerosis.^21, 22^ Of note, we have reported that a similar kinase inhibitor, “sorafenib,” is cardiotoxic, despite an anti-fibrotic effect.^22^ Paradoxically, our study demonstrated the cardioprotective effects of NTB in a preclinical model of HF. In fact, with our multiple years of experience researching the cardiac effects of various kinase inhibitors, particularly cancer therapeutic KIs, NTB is the first with favorable outcomes at both fronts, cancer, and heart.

Since we observed a reduction in cardiac hypertrophy in NTB treated animals, we were interested in determining if NTB directly affects cardiomyocytes apart from its well-studied fibroblast specific anti-fibrotic mechanisms. Although NTB could not prevent agonist-induced CMs hypertrophic response, significant alteration in key signaling pathways underlying such stimulus was observed. Of note, the expression and biological function of NTB targeted receptors in CMs are well-established. Therefore, it was not surprising to observe a significant effect of NTB on major cardiomyocyte signaling pathways. Taken together, it is conceivable that NTB’s cardioprotective effects can not be simply attributed to its direct impact on fibroblast and fibrosis alone. Further studies are warranted to delineate detailed mechanisms of NTB’s therapeutic efficacy and define the contribution of various cells, including fibroblast and cardiomyocytes.

Noth et al. investigated the cardiovascular safety of NTB in IPF patients.^23^ They perormed an analysis using pooled data from the TOMORROW and INPULSIS trials and concluded that irrespective of cardiovascular risk at baseline, the incidence of major adverse cardiac events was similar between patients treated with NTB vs placebo. However, the study had limitations such as the absence of randomization at baseline and unavailability of data on cardiovascular medication. Thus, there is a need to evaluate the efficacy of NTB specifically in CVD patients. Importantly, the current standard-of-care therapeutic agents, such as angiotensin II receptor blockers (ARBs), beta-blockers, angiotensin-converting enzyme (ACE) inhibitors, statins, etc. operate through pleiotropic mechanisms and none of them are explicitly designed to target cardiac fibrosis. Hence, one can test the hypothesis that there might be synergistic benefits of NTB when given as combinatorial regimens with standard cardiac therapeutic agents.

In conclusion, our study showed beneficial effects of NTB on pressure overload-induced cardiac remodeling and dysfunction in mice. We further demonstrated that the cardioprotective effects of NTB are mediated via multiple coordinated mechanisms directly affecting cardiac fibroblasts and cardiomyocytes. Our findings provide a proof of concept for the repurposing of NTB in cardiac diseases and suggest the need for randomized, placebo-controlled clinical studies to evaluate the safety and efficacy of NTB in HF patients.

## Sources of funding

This work was supported by research grants to HL from the NHLBI (R01HL143074-01A1, R01HL133290), PU was supported by American Heart Association (19POST34460025).

## Disclosures

None

## Notes

### Competing Interest Statement

The authors have declared no competing interest.

