## Supplemental Tables for "Repurposing Nintedanib for Pathological Cardiac Remodeling and Dysfunction"

Supplemental Table 1: Antibodies and dilutions used for Western blot analysis

| S.No. | Antibody | Vendor | Catalog No. | Dilution |
|---|---|---|---|---|
| 1 | p-ERK1/2 (Thr202/Tyr204) | Cell Signaling Technology | 4370S | 1:1000 |
| 2 | ERK1/2 | Santa Cruz Biotechnology | sc-93 | 1:1000 |
| 3 | p-AKT (Ser473) | Cell Signaling Technology | 4060 | 1:1000 |
| 4 | AKT | Cell Signaling Technology | 9272 | 1:1000 |
| 5 | p-p38 (Thr180/Tyr182) | Cell Signaling Technology | 9211s | 1:1000 |
| 6 | p38 | Santa Cruz Biotechnology | sc-535 | 1:1000 |
| 7 | pSMAD3 | Abcam | ab52903 | 1:1000 |
| 8 | SMAD2/3 | Cell Signaling Technology | 8685 | 1:1000 |
| 9 | pGSK-3α/β | Cell Signaling Technology | 9331 | 1:1000 |
| 10 | GSK-3α/β | Cell Signaling Technology | 5676 | 1:1000 |
| 11 | Fibronectin | Abcam | Ab2413 | 1:1000 |
| 12 | α-Tubulin | Santa Cruz Biotechnology | sc-8035 | 1:1000 |
| 13 | IRDye 680LT Goat anti-Rabbit IgG | LI-COR Biosciences | 926-68021 | 1:3000 |
| 14 | IRDye® 800CW Goat anti-Mouse IgG | LI-COR Biosciences | 925-32210 | 1:3000 |

Supplemental Table 2: TaqMan gene expression assays and control used for qPCR analysis

| **S.No.** | **Gene** | **Assay ID** | **Catalog No.** | **Vendor** |
|---|---|---|---|---|
| 1 | COL1A1 (Mouse) | Mm00801666_g1 | 4331182 | Applied Biosystems |
| 2 | COL3A1 (Mouse) | Mm00802300_m1 | 4331182 | Applied Biosystems |
| 3 | ANP (Rat) | Rn00664637_g1 | 4331182 | Applied Biosystems |
| 4 | BNP (Rat) | Rn00580641_m1 | 4331182 | Applied Biosystems |
| 5 | Eukaryotic 18S rRNA Endogenous Control | | 4319413E | Applied Biosystems |
